# Genetic patterns in *Montipora capitata* across an environmental mosaic in Kāne’ohe Bay

**DOI:** 10.1101/2021.10.07.463582

**Authors:** Carlo Caruso, Mariana Rocha de Souza, Lupita Ruiz-Jones, Dennis Conetta, Joshua Hancock, Callum Hobbs, Caroline Hobbs, Valerie Kahkejian, Rebecca Kitchen, Christian Marin, Stephen Monismith, Joshua Madin, Ruth Gates, Crawford Drury

**Affiliations:** Hawai’i Institute of Marine Biology, University of Hawai’i at Mānoa, Kāne’ohe, HI 96744, USA; Chaminade University of Honolulu, Honolulu, HI 96816, USA; University of Rhode Island, Kingston, RI 02881, USA; University of York, Heslington, York YO10 5DD, UK; Yale University, New Haven, CT 06520, USA; Stanford University, Stanford, CA 94305, USA

**Keywords:** *Montipora capitata*, Kāne’ohe Bay, Clonality, Genetic Relatedness, Seascape Genomics, environmental mosaic

## Abstract

Spatial genetic structure (SGS) is important to a population’s ability to adapt to environmental change. For species that reproduce both sexually and asexually, the relative contribution of each reproductive mode has important ecological and evolutionary implications because asexual reproduction can have a strong effect on SGS. Reef building corals reproduce sexually, but many species also propagate asexually under certain conditions. In order to understand SGS and the relative importance of reproductive mode across environmental gradients, we evaluated genetic relatedness in almost 600 colonies of *Montipora capitata* across 30 environmentally characterized sites in Kāne’ohe Bay, O’ahu, Hawai’i using low-depth restriction digest associated sequencing. Clonal colonies were relatively rare overall but influenced SGS. Clones were located significantly closer to one another spatially than average colonies and were more frequent on sites where wave energy was relatively high, suggesting a strong role of mechanical breakage in their formation. Excluding clones, we found no evidence of isolation by distance within sites or across the bay. Several environmental characteristics were significant predictors of the underlying genetic variation (including degree heating weeks, time spent above 30°C, depth, sedimentation rate and wave height); however, they only explained 5% of this genetic variation. Our results show that colony fragmentation contributes to the ecology of *M. capitata* at local scales and that genetic diversity is maintained despite strong environmental gradients in a highly impacted ecosystem, suggesting potential for broad adaptation or acclimatization in this population.

## Introduction

Coral reefs are diverse ecosystems that support a disproportionate number of the world’s marine organisms and provide valuable ecosystem services to humans. The health of reefs is determined by a complex interaction of environmental and biological factors, where coral assemblage structure is tightly linked to reef function. Despite their ecological importance, coral reefs are declining worldwide (De’ath, Fabricius, Sweatman, & Puotinen,2012; Hughes et al., 2017; Pandolfi et al., 2003). Major anthropogenic factors including climate change, pollution from terrestrial runoff, dredging, overfishing, and coastal development are negatively impacting corals, while interactions among multiple stressors often amplify declines (Muthukrishnan & Fong, 2014). Climate change induced heat stress can be exacerbated by local stressors, resulting in increased coral bleaching and mortality (Donovan et al., 2021; Hoegh-Guldberg, 1999; Sully, Burkepile, Donovan, Hodgson, & van Woesik, 2019; van Oppen & Lough,2018). But see Hughes et al., (2017). The decline of coral reefs has prompted a growing interest in developing active management solutions (van Oppen et al., 2017), which are dependent on the characterization of environmental and biological factors and their interactions.

Kāne’ohe Bay has a history of regular natural and anthropogenic disturbance; however, coral cover remains at ~60% (Bahr, Jokiel, & Toonen, 2015). The reefs in this ecosystem experience a broad range of conditions including diel variability in temperature, pH, and oxygen concentration (Barott et al., 2021; Drupp, De Carlo, Mackenzie, Bienfang, & Sabine, 2011;Drupp et al., 2013; Guadayol, Silbiger, Donahue, & Thomas, 2014; Shamberger et al., 2011) driven by the timing of low tide, water residence times and flow dynamics (Koweek et al., 2015;Lowe, Falter, Monismith, & Atkinson, 2009a). These interacting physical and biochemical processes create small-scale variation over as little as 25m(Guadayol, Silbiger, Donahue, & Thomas, 2014). Environmental differences over large and small spatial scales can generate intraspecific genetic divergence in corals, which may represent local adaptation to salinity, water chemistry, sedimentation and temperature (Bay & Palumbi, 2014; Cooke et al., 2020;Dixon et al., 2015; Selmoni et al., 2021; Selmoni, Rochat, Lecellier, Berteaux-Lecellier, & Joost,2020). Genetic divergence also can be driven by local anthropogenic pressure (Zvuloni et al.,2008), where corals in similar conditions tend to be more genetically similar (Tisthammer, Forsman, Toonen, & Richmond, 2020).

Clonal propagation occurs in many coral species, including branching and massive morphologies across a range of environments, suggesting the ubiquity of this process as an alternative reproductive strategy (Adjeroud et al., 2014; Baums, Miller, & Hellberg, 2006; Drury, Greer, Baums, Gintert, & Lirman, 2019; Foster et al., 2013; Gélin et al., 2017; Gorospe & Karl,2013; Manzello et al., 2019). The dominant coral species in Kāne’ohe Bay, *Porites compressa* and *Montipora capitata* (Bahr, Jokiel, & Toonen, 2015), are capable of propagating asexually and demonstrate variable levels of clonality (Hunter, 1993; Locatelli & Drew, 2019; Nishikawa, Kinzie, & Sakai, 2009). Previous work in Kāne’ohe Bay shows that clonality ranges from low (Genet:Ramet <0.05) (Locatelli & Drew, 2019) to moderate (G:R ~0.5) (Nishikawa, Kinzie, & Sakai, 2009) levels in *M. capitata*, despite the existence of substantial fragmentation potential (Jokiel, Hildemann, & Bigger, 1983) and high survival rates of fragments (Cox, 1992).

The aim of this study was to examine the genetic relatedness of *Montipora capitata* across an environmental mosaic in order to understand spatial patterns of clonality and genetic structure. To do so, we mapped and sampled tissue from approximately 600 colonies from 30 sites at which we also measured environmental characteristics (temperature, sedimentation rates, and wave energy). This study also serves as a foundation for long-term monitoring of a model coral population in context of environmental patterns and processes in Kāne’ohe Bay.

## Methods

### Site and Colony Selection

In 2017, we established long-term study sites at 30 patch reefs spanning 12 km across Kāne’ohe Bay, O’ahu, Hawai’i (Figure 1a; Figure S1). The bay was divided into five blocks from South to North based on the water flow regimes and modeled water residence time (Lowe, Falter, Monismith, & Atkinson, 2009a; b). Block 1 has the longest water residence time with >30 days on average, blocks 2 and 3 have a residency of 10-20 days, and blocks 4 and 5 have typical residency times less than 1 day. We defined patch reefs in QGIS (QGIS Development Team, 2017) based on benthic habitat maps (Hawai’i Statewide GIS Program, 2017; Neilson, Blogett, Gewecke, Stubbs, & Tejchma, 2014) and imagery available in the OpenLayers plugin. Features were considered patch reefs if they were distinct coral reef structures not contiguous with coral structure of the fringing reef or fore-reef. All patch reef polygons were assigned to a block. For each block, 30 random GPS coordinates were generated within the patch reef polygons using random points function in QGIS. Coordinates that fell on the patch reef including Moku o Lo’e Island (Hawai’i Institute of Marine Biology) in block 2 were excluded from site selection because of the terrestrial influence of the island and the significantly altered reef ecosystem due to research.

**Figure 1.**
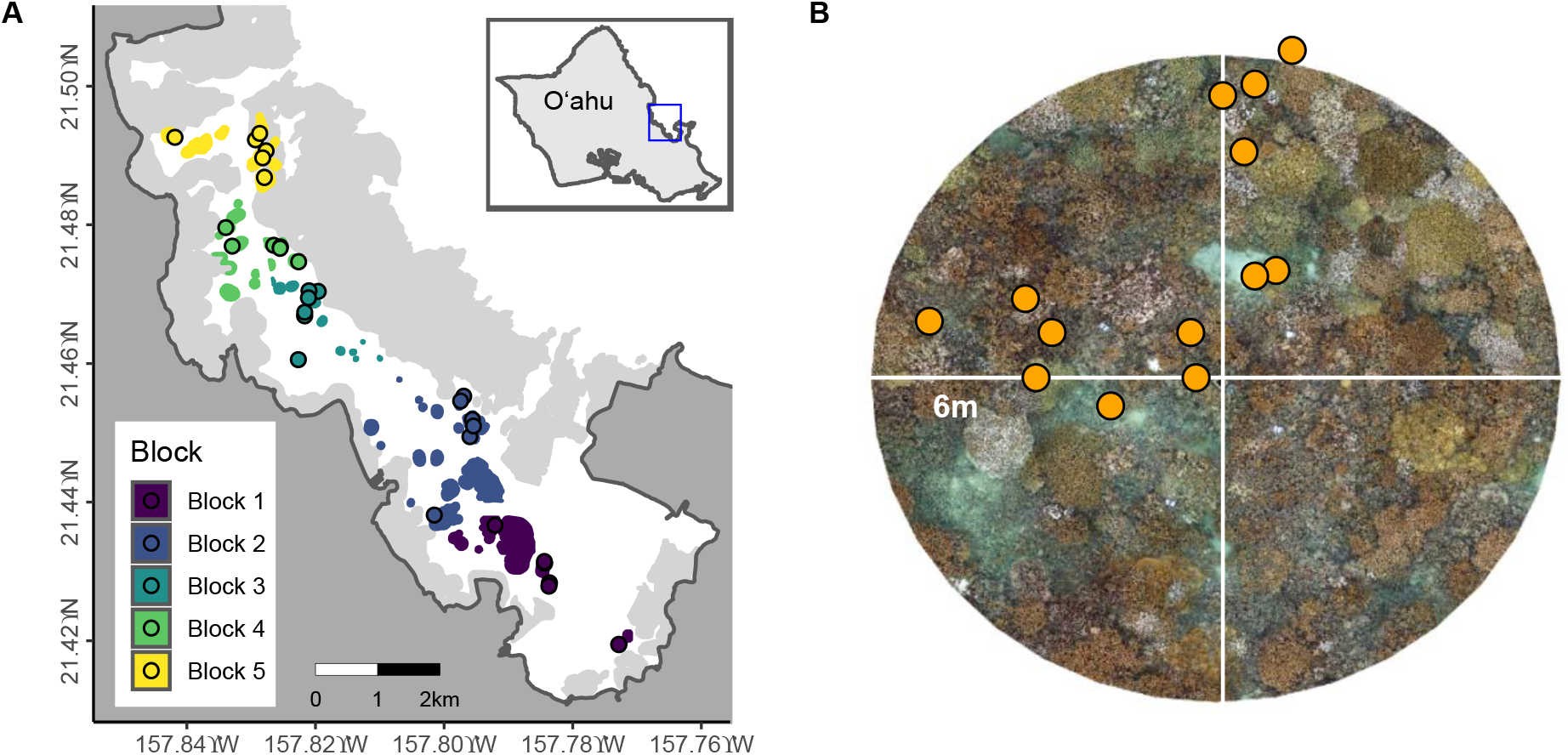
Study System. A) Map of Kāne’ohe Bay with colors denoting 30 sites distributed across 5 blocks. Sites were chosen using a random stratified design on patch reefs and approximately 20 *M. capitata* colonies were sampled from each. B) Example of an individual site, where corals are selected along a 10 m transect running in up to 4 cardinal directions from the center. Photomosaic background for example only, corresponding to 6m radius from central cinder block.

The random points were surveyed in sequential order and the first six suitable sites in each block were retained for the study, with a total of 30 sites across the bay (Figure 1a; Table S1). Sites were considered suitable if at least twenty colonies of *Montipora capitata* > 10 cm diameter were found in the initial survey and if the site was confirmed to be on a patch reef. Coordinates that fell very close to the edge of a patch reef were excluded. For selected sites, the central coordinate point was marked with a cinder block and nominal depth was recorded. Starting at the cinder block, a 10 m transect tape was laid consecutively to the west, south, east, and north directions until twenty colonies of >10cm diameter were found (**Figure 1B**; <10m from central block). Only corals that appeared visually healthy and thus physiologically and reproductively recovered from the 2014/2015 bleaching event (Cunning, Ritson-Williams, & Gates, 2016; Johnston, Counsell, Sale, Burgess, & Toonen, 2020; Ritson-Williams & Gates, 2020;Wall, 2019; Wall, Ritson-Williams, Popp, & Gates, 2019) were chosen for this study and individually tagged. Each colony was photographed with a color card standard and size scale after tagging. The position of the colonies along the transect was recorded, which later allowed us to calculate the distance between colonies within the same site.

Sites were periodically revisited, tags cleaned of biofouling, and colonies re-photographed and scored for health. During early 2019, colonies were individually assessed and ambiguous colonies that could no longer be definitely associated with a tag (either because the tag was lost, or the colony died) were noted and excluded from subsequent monitoring. Remaining colonies were re-tagged.

### Environmental Monitoring

Temperature data was recorded at 10-minute intervals with calibrated Hobo Pendant or Water Temp Pro v2 loggers (Onset Computer Corp., Bourne, MA) attached to the cinder block at the center of each site. Deployments began in July 2017 with loggers periodically retrieved, downloaded, recalibrated and/or cross-validated, and replaced at approximately 6–9-month intervals. Temperature monitoring is ongoing.

Sediment traps (16” vertically mounted capped 2” PVC pipe) were attached to each cement block. Sediment traps were exchanged every 1-2 months for a total of 7 deployments and the dry weight of recovered material was used to estimate the sediment accumulation rate for each site following (Storlazzi, Field, & Bothner, 2011).

Current meters (TCM-x w/MAt1 data logger, Lowell Industries, North Falmouth, MA) were deployed at one location in each block from March 18 to March 29, 2019. Meters were anchored to a cement slab following manufacturer’s installation guidelines and used to calculate root mean square (RMS) water velocity and wave height, which was generalized to nearby sites within the block.

### DNA Sampling, ddRAD Library Preparation, and Sequencing

A <1cm^2^ fragment was sampled in early 2018 from each tagged colony, preserved in 70% ethanol, and stored at −20°C until processed. We extracted DNA from a small piece of each fragment using Nucleospin Tissue Kits (Macherey-Nagel, Düren, Germany) following manufacturer instructions and quantified by fluorimetry (Quant-it HS dsDNA kit, Thermo-Fisher). We followed the general strategy of double-digest restriction-site associated DNA sequencing (ddRAD) outlined in (Peterson, Weber, Kay, Fisher, & Hoekstra, 2012). Briefly, we used an *in silico* digestion (*ddradseqtools* Python package (Mora-Márquez, García-Olivares, Emerson, & López de Heredia, 2017)) of the *Montipora capitata* genome (Shumaker et al.,2019) to choose enzymes expected to yield ~3000 fragments in the 220-240bp range.

*BclI* and *EcoRI* restriction enzymes (New England Biolabs (NEB), Ipswich, MA) were used to digest ~300 ng DNA from each sample in 30 μL reactions. We then ligated adapters (Integrated DNA Technologies, Coralville, IA), which included sequences complementary to the restriction cut motifs and sites for annealing PCR primers, with T4 DNA ligase (NEB). Samples were amplified (Q5 High-Fidelity Polymerase Kit, NEB) with a unique pair of primers based on Illumina TruSeq sequences (Illumina Inc., San Diego, CA), each containing a custom 6 bp barcode and a variable/degenerate 4 bp sequence for detecting PCR duplicates, along with p5/p7 flanking primers to enhance production of full-length constructs. The constructs were quantified by fluorimetry as above, reduced into 12 sub-pools of ~50 samples each, and size-selected on a Pippin Prep electrophoresis recovery instrument (Sage Science, Beverly. MA) using “Tight” mode with a 370 bp target size. Each sub-pool was then re-quantified, and all sub-pools were combined into an equimolar final pool. Clean up with Ampure XP SPRI beads (Beckman Coulter, Indianapolis, IN) was performed after ligation, subpooling, and final pooling steps to remove reagent contamination and for rough size selection.

The final sequencing library was composed of 640 uniquely barcoded samples. Replicates (69 biological replicates from 10 colonies (n=6-7 per colony) with 2 colonies from each block) were included in the library to evaluate clonality following (Manzello et al., 2019). The library was sequenced on a single lane of an Illumina HiSeq 4000 using paired-end 150bp chemistry (GeneWiz, South Plainfield, NJ, USA).

### Environmental Data Analysis

There were gaps in temperature data throughout the time series due to lost or corrupted loggers. We used a machine learning approach to impute raw temperature data using the *missForest* package (Stekhoven & Bühlmann, 2011). After filtering raw temperature data to timepoints with records for >70% of sites (at least 21 sites), we imputed missing information with mtry=100 and ntree=100. This approach presents a uniformly incomplete time series from July 2017 to March 2019 by filling in 12.8% of the overall dataset (**Figure 2A**). We performed a principal component analysis of the time series in each block (**Figure 2B**).

**Figure 2.**
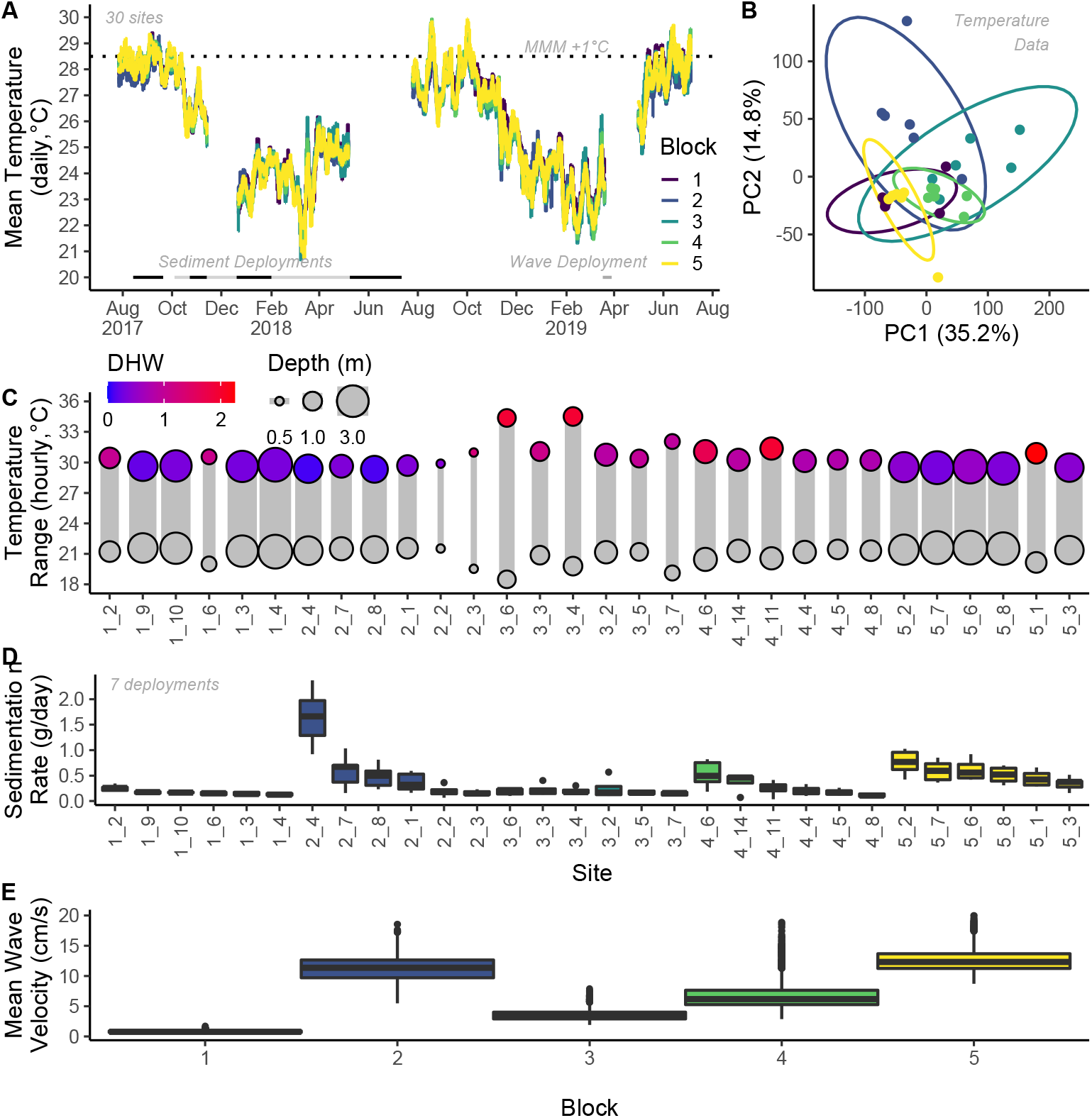
Environmental Characteristics. A) Temperature profiles from August 2017 to August 2019 for the 30 sites, color coded by block. B) PCA of all temperature data in panel A, color coded by block. Ellipses are 95% confidence intervals. C) Minimum and maximum hourly temperature at all sites. Gray bar width and point size correspond to nominal site depth and color corresponds to DHW accumulated over three summers (2017-2019). C) Mean sedimentation from 7 timepoints on square root transformed y-axis for visualization. E) Mean wave velocity in each block. Boxplots represent mean +/- 1 IQR.

From these data, we summarized hourly means and calculated DHW as time spent above 28.5°C (MMM + 1°C; (Dilworth, Caruso, Kahkejian, Baker, & Drury, 2021)) and summarised the total DHW through summers of 2017-2019. We also calculated the global mean, maximum and minimum temperature at each site and the average daily range experienced. We took the bay-wide average temperature for each day and calculated the residual, which was averaged for each site for the entire time period and calculated the total number of hours spent above 30°C. We used hourly averaged NOAA NCRMP temperature data from 2008 to 2019 at five nearshore O’ahu sites (<10m; (Pacific Islands Fisheries Science Center, 2021)) to evaluate local seasonality and defined the warmest stable period of the year (hereafter ‘summer’) as August 15 to October 15 (**Figure S2**). We calculated daily temperature profiles for each site during this period (**Figure S3**). Finally, we extracted data from the summer and calculated the three-year average for mean, standard deviation, daily range and minimum temperature. We averaged depth-corrected wave height, wave velocity (RMS), and sediment to produce inputs for the distance-based redundancy analysis (dbRDA).

### Analysis of genomic data

We trimmed adapters and removed PCR duplicates from sequencing data using tagseq_clipper.pl (https://github.com/z0on/tag-based_RNAseq) and then trimmed reads using Trimmomatic 0.39 (Bolger, Lohse, & Usadel, 2014) when the 5bp average quality score was <20. Reads were aligned to the *M. capitata* genome using bowtie2 1.3.0 (Langmead & Salzberg,2012) with --very-sensitive-local settings.

Aligned reads were processed to assess pairwise genetic distance using ANGSD 0.931 (Korneliussen, Albrechtsen, & Nielsen, 2014) with the following specifications: -doIBS 1, -minInd 320, -minQ 30, -minMapQ 20, -SNPpval 1e-4. We also calculated genotype likelihoods using ANGSD with allele frequency priors (-GL 2, -doGeno 8, -doPost 1, -minInd 200, -minQ 30, -minMapQ 20, -genoMinDepth 3). We then summed the probability of the heterozygote (ab) and 2 × secondary homozygote (bb) to predict the number of secondary alleles without hard-calling genotypes for each sample following (Drury & Lirman, 2021).

We used the relatedness from ANGSD to calculate the 95th percentile of pairwise distance between biological replicates after visually assessing outliers (**Figure 3A**); this value was used as the threshold for calling genotypes using hierarchical clustering with the complete method in the R package *hclust*. After determining the existence of clonal groups, we calculated the Genet-Ramet ratio by dividing the number of genotypes by the number of samples at each site and calculated mean relatedness with and without clones at each site. We used a Wilcoxon test to compare the spatial distance of all non-clonal samples at a site with the distance of clonal samples to evaluate spatial clustering. We used a Mantel test to calculate isolation by distance after randomly selecting one individual from each genotype at each site using the R package *ade4* (Dray & Dufour, 2007). Wilcoxon and mantel p-values were FDR adjusted to account for multiple comparisons. To test for the effect of physical disturbance, we used a linear regression to compare genet:ramet ratio to wave height and depth. We also used a Mantel test to compare geographic and genetic distance across Kāne’ohe Bay. We processed genotype likelihood values as described above as the summary statistic of the genotype of each sample at each locus and randomly excluded all but one sample from each genotype to remove clonality from the downstream analysis. We used a distance based redundancy analysis (dbRDA) with Bray-Curtis distances to project genetic variation between individuals explained by environmental factors into reduced dimensional space (Capblancq, Luu, Blum, & Bazin,2018), with an ANOVA to test for overall significance using the R package *vegan* (*Dixon, 2003*). We used a PERMANOVA to test for significant differences between blocks and calculated variance explained using sum of squares of all environmental characteristics. All statistical analyses were performed in R 4.0.4 (R Core Team 2021)

**Figure 3.**
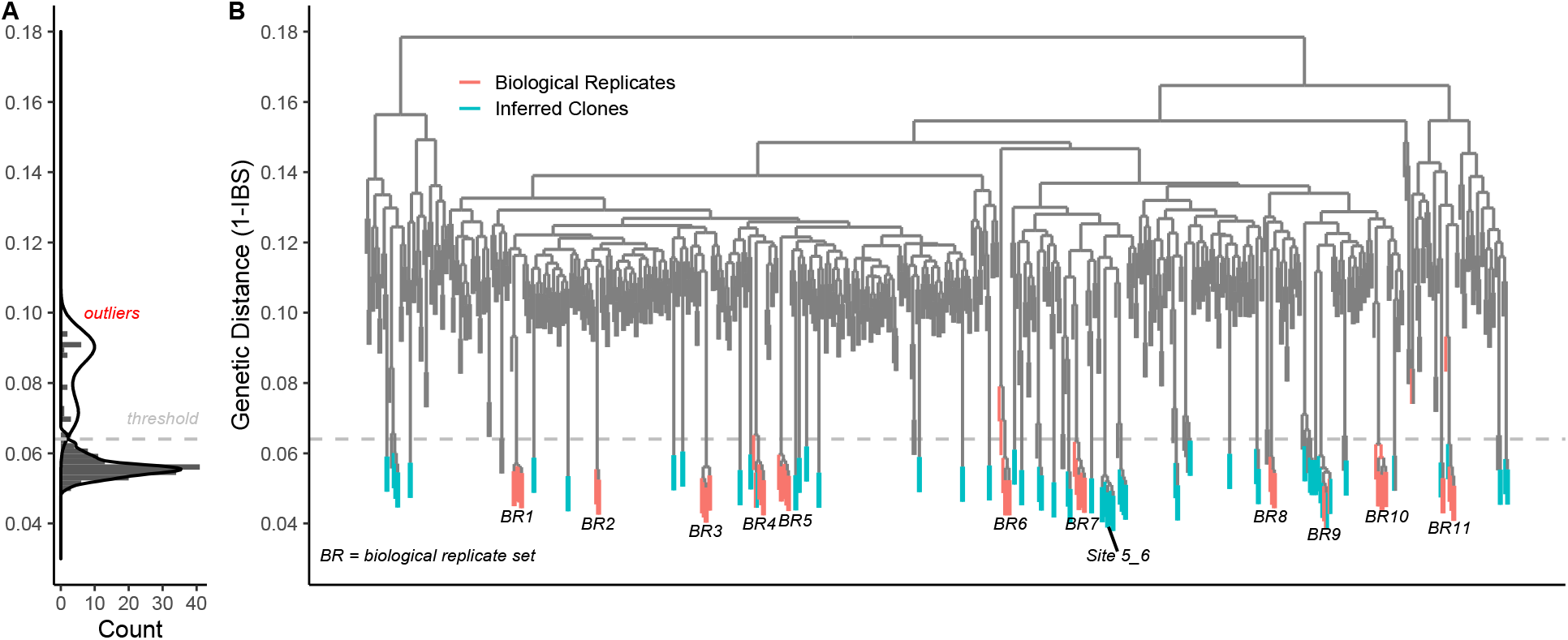
Patterns of Clonality in *M. capitata* across Kāne’ohe Bay. A) Distribution of pairwise genetic distances in biological replicates (69). Red highlighted values were visually inspected and designated as outliers. Gray values represent distribution from which 95th percentile was calculated to determine clones in the broader population. B) Identity by descent dendrogram calculated using complete hierarchical clustering. Orange groupings represent biological replicates (n=10) and blue groupings represent inferred clones. One genotype at site 5_6 was composed of 8 colonies, but most other clones were pairs of colonies within the same site.

## Results

The 30 sites in this study had broadly different environmental characteristics (**Figure 2**, **Supplemental Table 1**). Temperatures followed the same seasonal trajectory throughout the bay (**Figure 2A**) but had substantially different fine-scale dynamics including minimum, maximum, and daily range (**Figure 2C**). During the 2.5 years, corals experienced between 0 and 2.2 DHW (**Figure 2C**) but had substantially different daily ranges (0.51 to 2.58°C), including peak temperatures >34°C at 2 sites (3_4 and 3_6). Reefs experienced between 0 and 148 hours above 30°C during the 2.5 years. Nominal depth ranged from 0.5 to 3.5m (**Figure 2C**), sediment capture ranged nearly 300-fold from 0.01 g/day to 2.93 g/day (**Figure 2D**), wave height ranged from 0.08 to 0.31m (**Figure 2E**), and mean water velocity ranged 40-fold from 0.8 to 12.56 cm/s (**Figure 2**, **Supplemental Table 1**).

After filtration, QC and alignment, samples had a mean of 209,462 ± 136,798 reads (mean ± SD). We used ANGSD for this analysis, which is suitable for low and variable depth sequencing. The genetic distance (1-IBS) of biological replicates from the same colony was between 0.04 and 0.15; however, there was a clear set of outliers >0.07 (**Figure 3A**). The 95th percentile of the non-outlier pairwise distances was 0.064, which was used as the clonality threshold following (Drury, Greer, Baums, Gintert, & Lirman, 2019) (**Figure 3**).

All but one biological replicate of each sample was excluded resulting in a total of 579 samples used in clonality analysis. These samples represented 531 genotypes, of which 36 were found to have clones for a bay-wide genet/ramet ratio of 0.917. The most abundant genotype was composed of 8 sampled colonies (**Figure 3B**), all of which were at a single site (5_6) in block 5, which had the lowest genet:ramet ratio of any site (G:R=0.578; **Supplemental Table 2**). Three genotypes included colonies from multiple reefs (n=2-3 reefs), all of which were in block 2, which has high wave energy (mean 11.178 rms). Most clonal genotypes contained only 2 representative colonies at the same site (26 of 36). G:R ranged from 0.578 to 1, and 11 sites had G:R equal to 1, indicating no clones were found (**Supplemental Table 2**).

We used dbRDA to examine environmental drivers of genetic variation and found eight factors were significant after multiple comparisons correction (p<0.05; **Figure 4A** and **Supplemental Table 3**). In order of explained variance, these were depth, hours above 30°C, degree heating weeks, mean sedimentation, summer daily temperature range, minimum temperature, wave height, and temperature residual. There was significant population structuring between blocks (**Figure 4B**; PERMANOVA p<0.001). We examined the presence of samples from corals that were members of a clonal genotype (only one colony from each genotype was included) in the dbRDA analysis and found that clonal colonies were similar to the overall distribution of genetic variance across the bay (**Figure 4C**).

**Figure 4.**
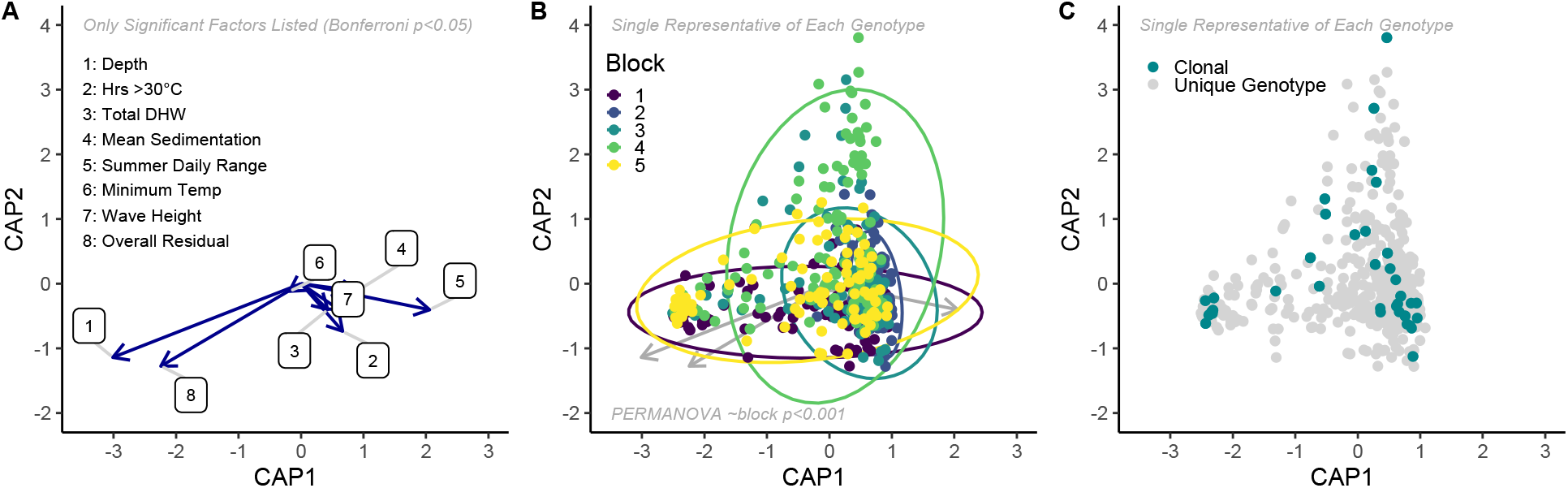
dbRDA analysis. Distance based redundancy analysis was used to examine environmental drivers of underlying genetic patterns. Points represent individual colonies in the study, with only one random individual from each clonal group (i.e. clones removed). A) Significant (Bonferroni p<0.05) environmental drivers of genetic patterns in the bay. Each arrow signifies the multiple partial correlation of the environmental driver in the RDA whose length and direction can be interpreted as indicative of its contribution to the explained variation. Environmental variables are listed in descending order of variance explained. B) Distribution of colonies based on genotype likelihoods, colored per block. Ellipses are 95% confidence intervals. C) Distribution of corals with clonal replicates in the bay. For all dbRDA analysis, only one individual sample from a genotype was used to avoid clonality effects in the analysis.

There was little variation in relatedness among sites, with mean values within a site ranging from 0.106 to 0.121 with clones included (**Figure 5BC**). When only one random sample from each clonal group at each site was included, the minimum mean relatedness increased to 0.114. Within sites there were no examples of significant isolation by distance after multiple comparisons correction (Mantel test FDR p>0.1), but at all sites with clones (n=19), colonies of the same genotype were significantly closer together than the average colony at that site (Wilcox test; FDR p<0.045). There was a significant negative relationship between G:R and wave height (lm p=0.047) and depth (lm p=0.046) which explained 31.4% of variance, indicating that wave energy is positively related to the presence of clones (lm p=0.123). We used a Mantel test to compare genetic distance (one instance of every genotype) and geographic distance over the entire population and found no relationship (**Figure S4**; R=-0.009, p=0.71).

**Figure 5.**
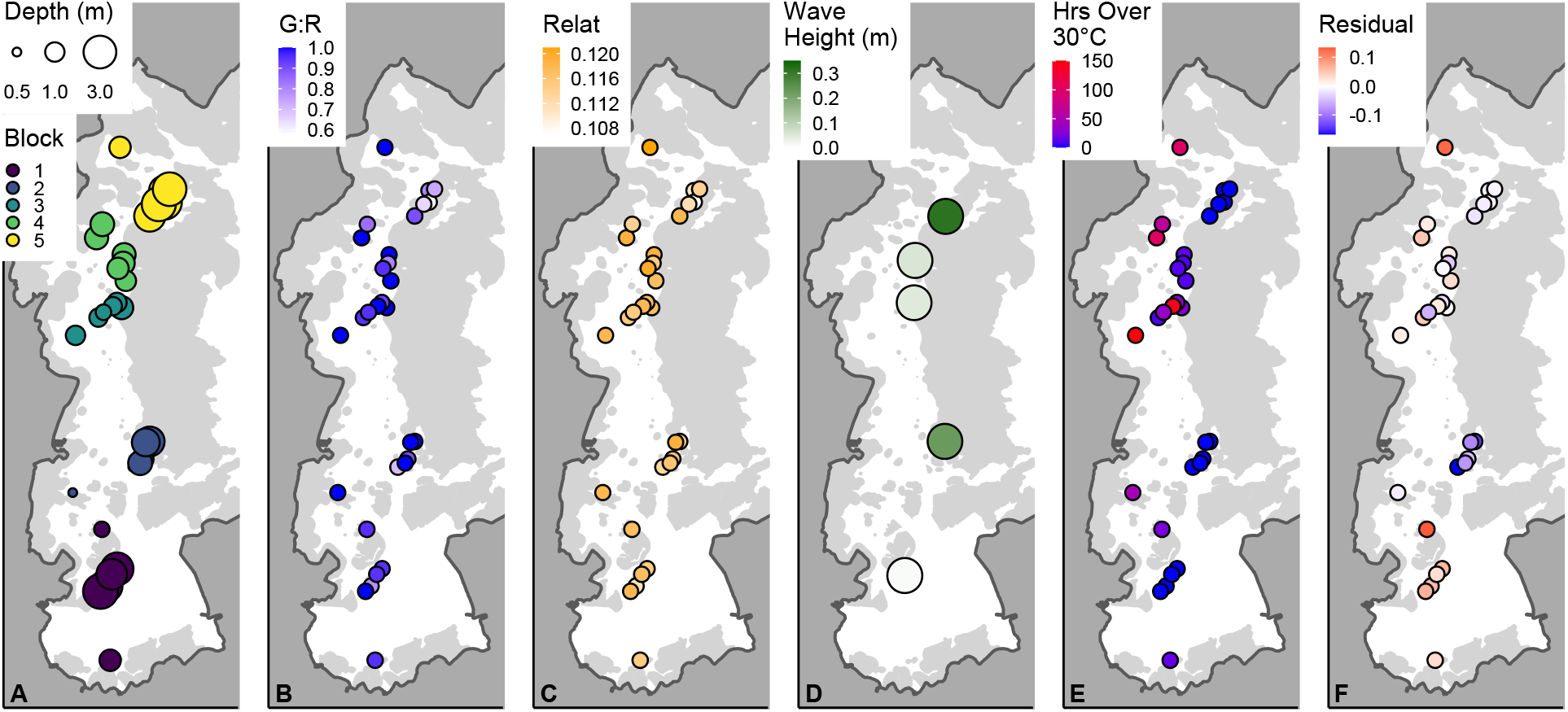
Spatial Distribution of Genetic and Environmental Outcomes. A) Map of Kāne’ohe Bay with colors denoting 5 blocks and circle size displaying depth. B) Genet:Ramet ratio at each site. C) Mean pairwise relatedness at each site, excluding all but one representative of each genotype. D) Mean wave height from a single site in each block. E) Hours over 30°C at each site. F) Average Residual temperature at each site.

Environmental characteristics of individual sites were highly variable across Kāne’ohe Bay (**Figure 5**). Wave energy was low in the South Bay (block 1) and at mid-bay sites shielded by the sandbar (**Figure 5D**; blocks 3,4). High temperature sites were concentrated in northern and inshore reefs (**Figure 5EF**), with negative residuals (cooler than average sites) centered in the relatively exposed block 2 (**Figure 5F**).

## Discussion

By investigating the genetic structure of almost 600 colonies of *Montipora capitata* across 30 sites in Kāne’ohe Bay, we show that clonality is rare and mostly restricted to habitats conducive to fragmentation by wave energy. We also show that non-clonal genetic relatedness across the bay is not correlated with distances among colonies, which indicates high levels of mixing and dispersal in sexually produced larvae. These results indicate a high potential for acclimation or adaptation of multiple genotypes to the environmental conditions present in the bay, rather than selection of a few more resilient ones.

Previous work has found strong associations of individual loci with environmental factors including temperature, salinity, wave action and water quality over spatial scales from 100s of meters to thousands of kilometers (Bay & Palumbi, 2014; Cooke et al., 2020; Jin et al.,2016; Selmoni, Rochat, Lecellier, Berteaux-Lecellier, & Joost, 2020; Selmoni et al., 2021). Population structure is also related to tidal flux (Underwood et al., 2020) and depth, although depth contrasts have primarily been between shallow and mesophotic reefs (Bongaerts et al.,2017; Serrano et al., 2014). Conversely, our study represents a range of sites across an environmental mosaic of depth, wave action, sedimentation, and temperature combinations in a semi-enclosed bay with a total distance separating sites of less than 12km. We show that no single environmental factor was disproportionately influential on genetic variation, but that depth, sedimentation rate, wave height and a number of temperature metrics were all significant (**Supplemental Table 3**). This suggests that tradeoffs may be important in an environmentally heterogeneous system, dampening strong local adaptation but creating some signals in the underlying genetic variation. In *M. capitata* in Kāne’ohe Bay, depth strongly dictates symbiosis state (Innis, Cunning, Ritson-Williams, Wall, & Gates, 2018), which covaries closely with host genetics (Drury et al., 2021). We hypothesize that light, which is strongly attenuated within the first few meters, acts as a selective pressure in this ecosystem via differential impacts on corals harboring *Cladocopium* and *Durusdinium*.

After depth and wave energy, the next most important explanatory variables were temperature characteristics of individual sites, including total degree heating weeks and hours >30°C, suggesting that some underlying genetic variation is associated with local conditions during the hottest time of the year. Interestingly, minimum temperature was also a significant predictor of this genetic variation, which may indicate tradeoffs between warm and cold tolerance (Howells, Berkelmans, van Oppen, Willis, & Bay, 2013) are an important factor for coral populations in Kāne’ohe Bay that experience large annual temperature variation of >15°C at some sites (this study).

We also found significant differences in genetic relatedness between blocks in Kāne’ohe Bay. Combined with the low variation explained by our dbRDA analysis, this is likely a signal of environmental influence outside of our measurements that relate to water residence times or other anthropogenic impacts that covary with gross position within Kāne’ohe Bay. There was a notable difference in genetic diversity between certain blocks; where block 4 varied along the second Principal Component (**Figure 4B**) which was not strongly related to any environmental variables described here, block 2 was particularly genetically constrained relative to other sites. Previous work across the Hawaiian archipelago, found that *M. capitata* from Kaneohe Bay was well connected with the other Main Hawiian islands sampled, including Maui and Hawai’i (Concepcion, Baums, & Toonen, 2014), despite distinctive higher anthropogenic pressure and warmer temperatures (Bahr, Jokiel, & Toonen, 2015). Locatelli & Drew (2019) also found signatures suggesting some genetic structure in *M. capitata* in Kāne’ohe Bay, although their sampling regime was more focused on inshore-offshore gradients, did not include the southern part of Kāne’ohe Bay, and likely did not capture the fine scale environmental mosaicism of our sites. This is likely the reason why we did not observe isolation by distance in our dataset, in keeping with the “coral population genetic paradox” observed in multiple coral studies (Gorospe & Karl, 2013).

Consistent with previous findings (Hunter, 1993; Jokiel, Hildemann, & Bigger, 1983;Nishikawa, Kinzie, & Sakai, 2009), we found that clonality is a significant but not a prominent feature of the *M. capitata* population of Kāne’ohe Bay patch reefs. For sites where we detected clonality, there was a clear spatial correlation with clonemates positioned more closely to one another relative to non-clonal colonies, consistent with expectations that genetic dissimilarity increases with distance. Clonality has been observed frequently within reefs, but the degree of clonality varies widely from almost none in some species-site combinations to nearly 100% in others (Hunter, 1993; Baums, Miller, & Hellberg, 2006; Drury, Greer, Baums, Gintert, & Lirman,2019; Foster et al., 2013; Jokiel, Hildemann, & Bigger, 1983; Manzello et al., 2019; Miller & Ayre, 2004).

Even within individual species, different patterns in clonality frequency have been observed across the species range, with some regions showing a higher prevalence of asexual recruitment that may be due to habitat differences (Adjeroud et al., 2014; Baums, Miller, & Hellberg, 2006). In other ecosystems, clonality is expected to be more prevalent in stable habitats which favor selected, reduced genetic diversity while habitats with more environmental variation or disturbance support more genetic diversity (see discussion in (Miller & Ayre, 2004)). In corals, this expectation may be complicated by the relationship between habitat stability and wave energy which is a major driver of the fragmentation mode of asexual reproduction. (Coffroth & Lasker, 1998) discussed expectations for fragmentation-based coral reproduction across physical disturbance levels and concluded that clonality will be highest where there is enough periodic physical disruption to promote some breakage but not enough to prevent survival and attachment of new propagules.

Fragmentation in corals could have adaptive value (Highsmith, 1982). Particular patterns of three-dimenionsality (Lasker, 1984) and increased growth (“pruning vigor”) following fragmentation in some species (Lirman et al., 2010) have been proposed as adaptations to enhance asexual propagation, although this is difficult to evaluate experimentally. We found no indication of a genetic basis for the frequency of fragmentation (**Figure 4C**) in our survey. Corals that were documented to be part of a multi-colony clone were distributed as expected in multivariate genetic space similarly to all other colonies in the site (some of which could also be clonal but were not sampled) indicating that no genetic variants relate to fragmentation. While genetic correlates with skeletal density and morphology may exist, our results support physical drivers of clonality; genet:ramet is lower (more clones) at sites with larger wave height consistent with stochastic fragmentation.

While our results show increased clonality at sites that are subjected to more wave energy, corals were only sampled at patch reefs which excluded habitats subjected to the highest levels of wave energy. Our sites with the most exposure to the ocean were the block 5 sites adjacent to the northern channel (mean G:R 0.73). Hunter (1993) included exposed fore reef sites outside Kāne’ohe Bay and observed a reduction in *Porites compressa* clonality both there (G:R 0.87) and in the more sheltered South Kaneohe Bay (SKB) sites (G:R 0.96) relative to North Kaneohe Bay (NKB) sites (G:R 0.64). SKB is in the vicinity of our sites 1_3 and 1_4 where we measured a mean G:R 0.89 and NKB is close to several sites in our block 4 where we measured a mean G:R of 0.92. These observations are consistent with the (Coffroth & Lasker,1998) predictions, indicating that some of the more exposed patch reefs in Kāne’ohe Bay may represent an optimal zone for *M. capitata* and *P. compressa* clonality.

Within a gradient of physical disturbance, substrate is also an important determinant of the outcomes of fragmentation (Coffroth & Lasker, 1998). In Kāne’ohe Bay, (Nishikawa, Kinzie, & Sakai, 2009) noted that the sheltered site (SHEL; equivalent to our site 1_2) had many small unattached colonies whereas the exposed site (EXPO; in vicinity of our sites 2_1, 2_2, 2_4, 2_7, 2_8) had mostly large attached colonies and that the ratio of genotypes to colonies was significantly lower for unattached versus attached colonies at SHEL, which is primarily sandy mud unconducive to fragment attachment. We found high genetic diversity at the equivalent sites (G:R for site 1_2 was 0.95 and mean G:R for sites at block 2 was 0.92); however, we did not include small unattached colonies in our sampling scheme. Inclusion of such colonies may have increased the level of clonality detected at sites with sufficient wave energy to fragment existing colonies, but insufficient hard substrate for them to establish themselves as fixed structures.

Jokiel, Hildemann, & Bigger (1983) examined *Montipora verrucosa* (presumed to be equivalent to what is presently identified as *Montipora capitata*) and *Montipora dilatata*, which is no longer common in Kāne’ohe Bay. These studies observed that patches of *M.dilatata* were greatly reliant on asexual reproduction (100% of colonies were histocompatible) while *M. verrucosa* patches were dependent on sexual reproduction (only 5% histocompatible). These results highlight wide intrageneric variability in clonal structure and the potential role of morphology in these patterns. *M. dilatata* has a thin branching structure and can easily be broken, which contrasts with the typically more substantial branches or plates of *M. capitata* and may explain the relative differences between species.

Corals have different modes of asexual reproduction including production of ameiotic parthenogenetic larvae, budding, and physical fragmentation, which interact with habitat disturbance to impact clonality patterns. For example, *Pocillopora acuta* in Kāne’ohe Bay release parthenogenetic larvae with moderate dispersal potential (comparatively lower than broadcast spawning larvae, but higher than fragmentation), achieving very high levels of clonality over the same spatial scale and in the same patch reef environments where we observed *M. capitata* (Gorospe & Karl, 2013). *P. acuta* in the Philippines can dominate reefs via asexual reproduction of ameiotic larvae, where clonality rates are highest with less wave energy (Torres, Forsman, & Ravago-Gotanco, 2020), but other studies have found no evidence of clonality regardless of habitat (Miller & Ayre, 2004). Another coral found in Kāne’ohe Bay, *Lobactis scutaria* (formerly *Fungia scutaria*) can form clones by budding and is often found in dense localized aggregations presumed to be largely derived from asexual reproduction (Lacks,2000).Conversely, *M. capitata* is a hermaphroditic broadcast spawner with rare self-fertilization (Padilla-Gamiño, Weatherby, Waller, & Gates, 2011) that is not known to produce parthenogenetic larvae. In an environment like Kāne’ohe Bay, this effectively restricts asexual propagules to the same patch reef where they are derived and supports our conclusions that patterns of clonality are driven by fragmentation. However, we did find three instances of colonies identified as the same genotype on separate patch reefs, all in block 2. Two of these instances were on neighboring reefs separated by ~300m, but the third comparison (2_3 and 2_4) is nearly 2km away. This is likely due to a sequencing error as there is no realistic way for fragmentation across the deeper portions of the bay to bridge this gap.

Clonality by fragmentation, budding, or production of asexual planula may be an adaptation to unfavorable environmental conditions (Foster et al., 2013). It provides advantage to a genet by increasing its long-term survival and reproductive success, allows species and genets to persist when unable to sexually reproduce (Honnay & Bossuyt, 2005) and enables well adapted genotypes to become locally dominant (Drury, Greer, Baums, Gintert, & Lirman,2019). For most scleractinian species, asexual reproduction is a result of fragmentation, where wave action can spread ramets over large distances, increasing the frequency and range of a genotype. Fragments may have higher chances of survival than planula because they surpass size specific mortality thresholds (Jackson, 1977) and may allow the colonization of areas on the reef not suitable for larval settlement (Highsmith, 1982). Furthermore, spawning in most corals is restricted to one or few months (Richmond & Hunter, 1990), while fragmentation can happen year-round. However, fragmentation limits the genetic diversity of populations not undergoing recombination during sexual reproduction and severe fragmentation can negatively impact corals, due to the energetic cost of lesion recovery, higher chances of infection and decreased fecundity due to size reduction (Zakai, Levy, & Chadwick-Furman, 2000; Lirman, 2000; Lirman et al., 2010; Smith & Hughes, 1998).

While clonality provides advantages to individual genets, high levels of genetic diversity within populations enhance ecosystem recovery after extreme events and is key for the longevity of those populations in a rapidly changing climate (Baums, 2008; Baums et al., 2019;Booy, Hendriks, Smulders, Groenendael, & Vosman, 2000; DiBattista, 2008). The decline of coral reefs has led to an increased interest in coral restoration efforts worldwide (reviewed in (Boström-Einarsson et al., 2020), but one concern is that the newly restored population will have less genetic diversity than the original population due to few donor colonies or small or non-existent genetic diversity among the donor colonies (Baums, 2008; Baums et al., 2019;Shearer, Porto, & Zubillaga, 2009), leading to genetic swamping or maladaptation. Incorporating metrics of genetic diversity in coral reef restoration efforts can provide valuable information when selecting coral colonies for restoration and can increase the effectiveness of restoration that may prove to be critical for the future or reefs.

Kāne’ohe Bay has been exposed to decades of conditions considered deadly for most corals worldwide (Bahr, Jokiel, & Toonen, 2015). The cessation of some point source stressors in the mid-1900s led to an increase in coral cover, but the bay still experiences heavy fishing pressure and higher temperatures and acidification than nearby reefs (Guadayol, Silbiger, Donahue, & Thomas, 2014; Jury & Toonen, 2019). Here we confirm high genetic diversity of *M. capitata* in Kāne’ohe Bay, finding few clones produced mostly by fragmentation in high wave energy sites. This outcome suggests that corals in this ecosystem have the potential for broad adaptation or acclimatization to this multistressor system and do not rely on selection and asexual propagation of a few resilient genotypes to recover and sustain functional ecosystems. Future studies incorporating bleaching dynamics, symbiont diversity, environmental variability and clonality would better help elucidate the selective pressures this species is undergoing and help predict how it will respond to the environmental conditions expected with climate change.

## Author Contributions

**LRJ, CC, MRDS**, and **RG** conceived the study. **CC, MRDS, LRJ, DC, JH, CH, CH, VK, RK, CM, SM** collected data.**CC**, **CD, CH, LRJ** analyzed data. **CC, MRDS, LRJ** and **CD** wrote the manuscript. **CD**, **RG**, **JM** and **SM** contributed funding and reagents. All authors edited the manuscript and approved the final version.

## Acknowledgements

We thank Carly Kenkel and Sarah Davies for supplying barcode adapters and Rob Toonen and Misha Matz for discussions on library preparation and sequencing approaches. We thank the Gates Coral Lab and Coral Resilience Lab for assistance with field and lab work. CC, FD, MRDS, and JH acknowledge the Paul G. Allen Family Foundation. MRDS acknowledges the American Association for University Women. LRJ is grateful for the financial support for travel provided by the Thinking Matters Fellowship at Stanford University. We dedicate this paper to our coauthor Ruth Gates who encouraged us to search for solutions to the coral reef crisis. Colonies were tagged and sampled under DAR Permit SAP 2018-03 to HIMB. This is HIMB contribution #xx and SOEST contribution #xx.

