## Supplemental Figures for "Genetic patterns in *Montipora capitata* across an environmental mosaic in Kāne’ohe Bay"

**Supplemental material**

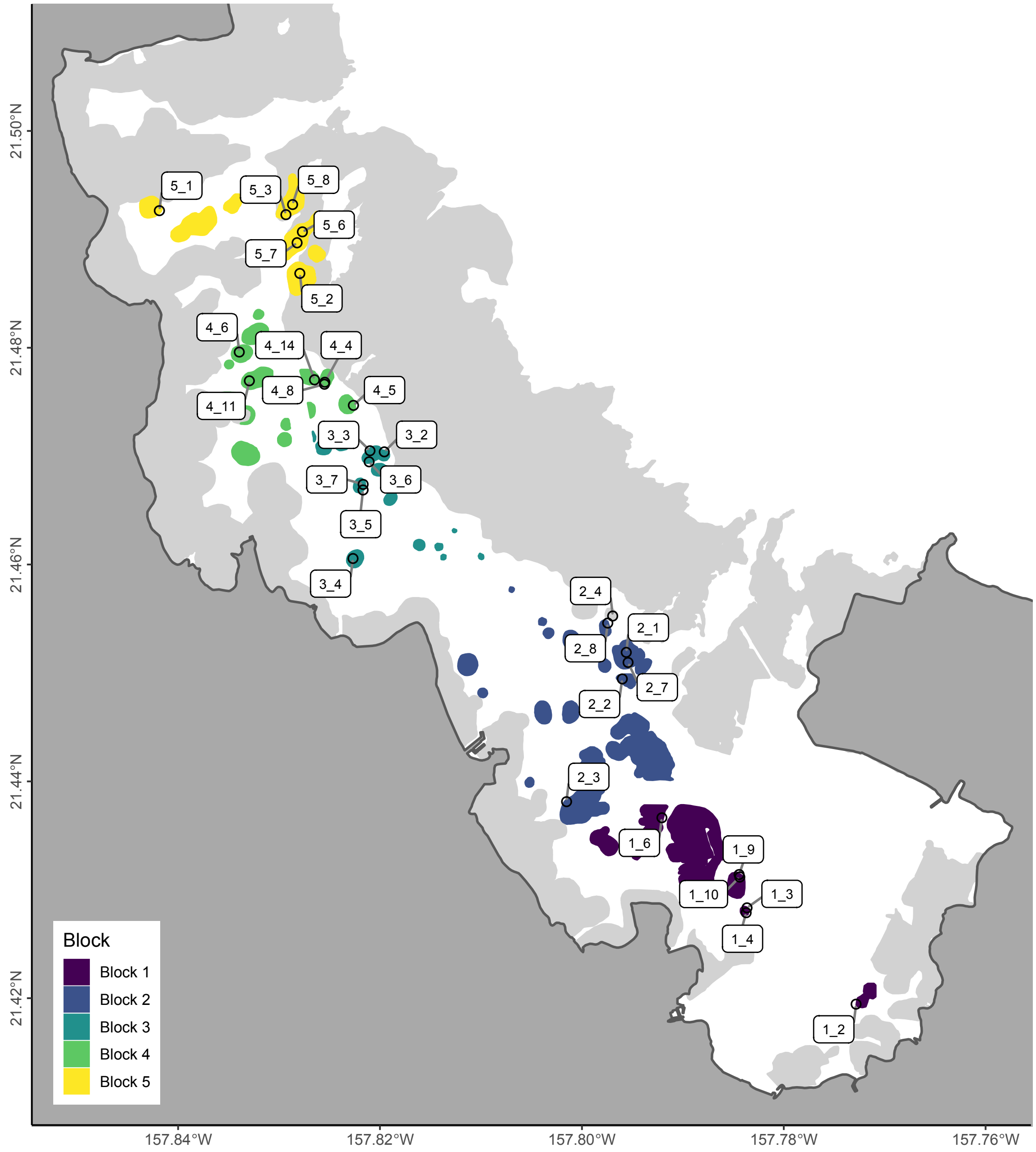


**Supplemental Figure 1 - Map of Study Sites**
Study sites, color coded by block with site names.

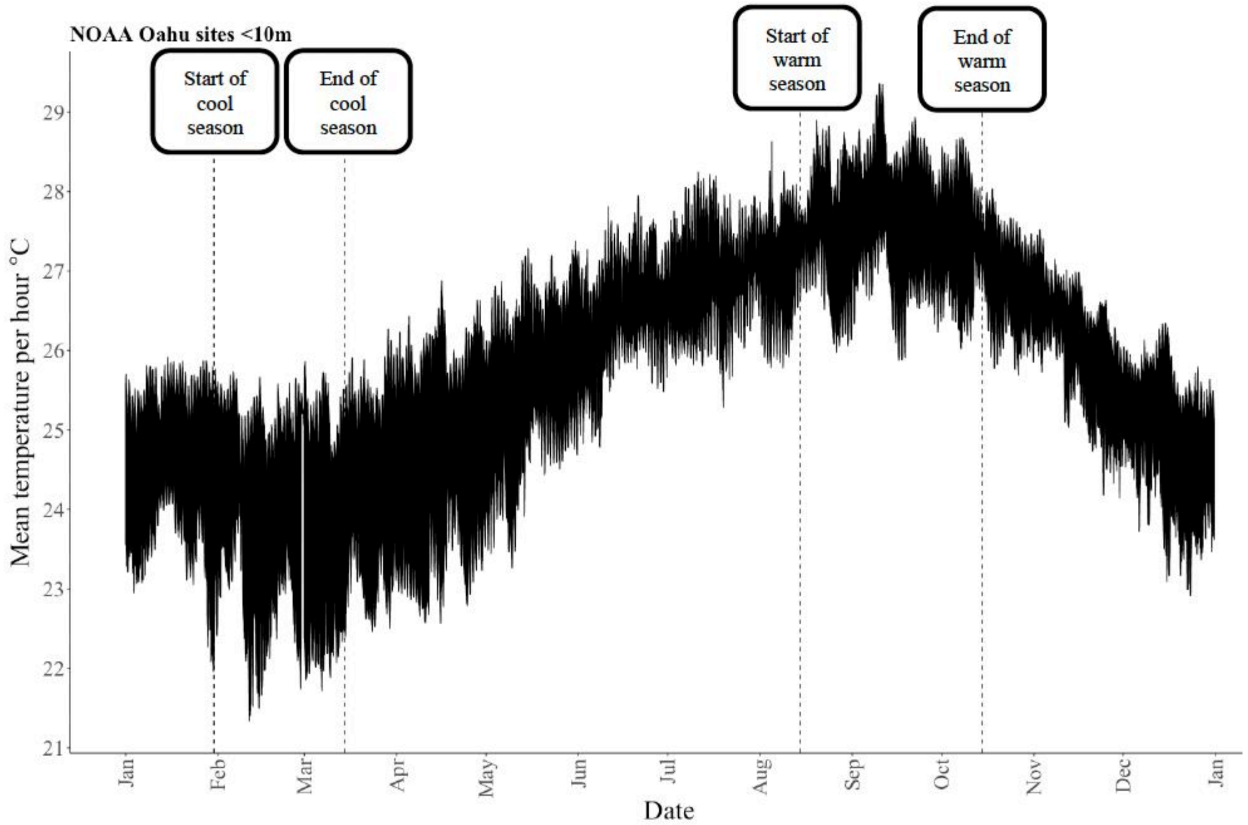


Fig 4. Seasonality of hourly mean water temperature (°C) at nearshore Oahu sites < 10 m in depth from 2008 to 2019. Temperature data were recorded using subsurface temperature recorders (STRs). Data taken from https://www.fisheries.noaa.gov/inport/item/37070.

**Supplemental Figure 2 - Water Temperature Seasonality on Oʻahu**

SST data from 2008 to 2019 for 5 sites <10m surrounding Oʻahu used to define the warm season for temperature calculations. Data from NOAA NCRMP.

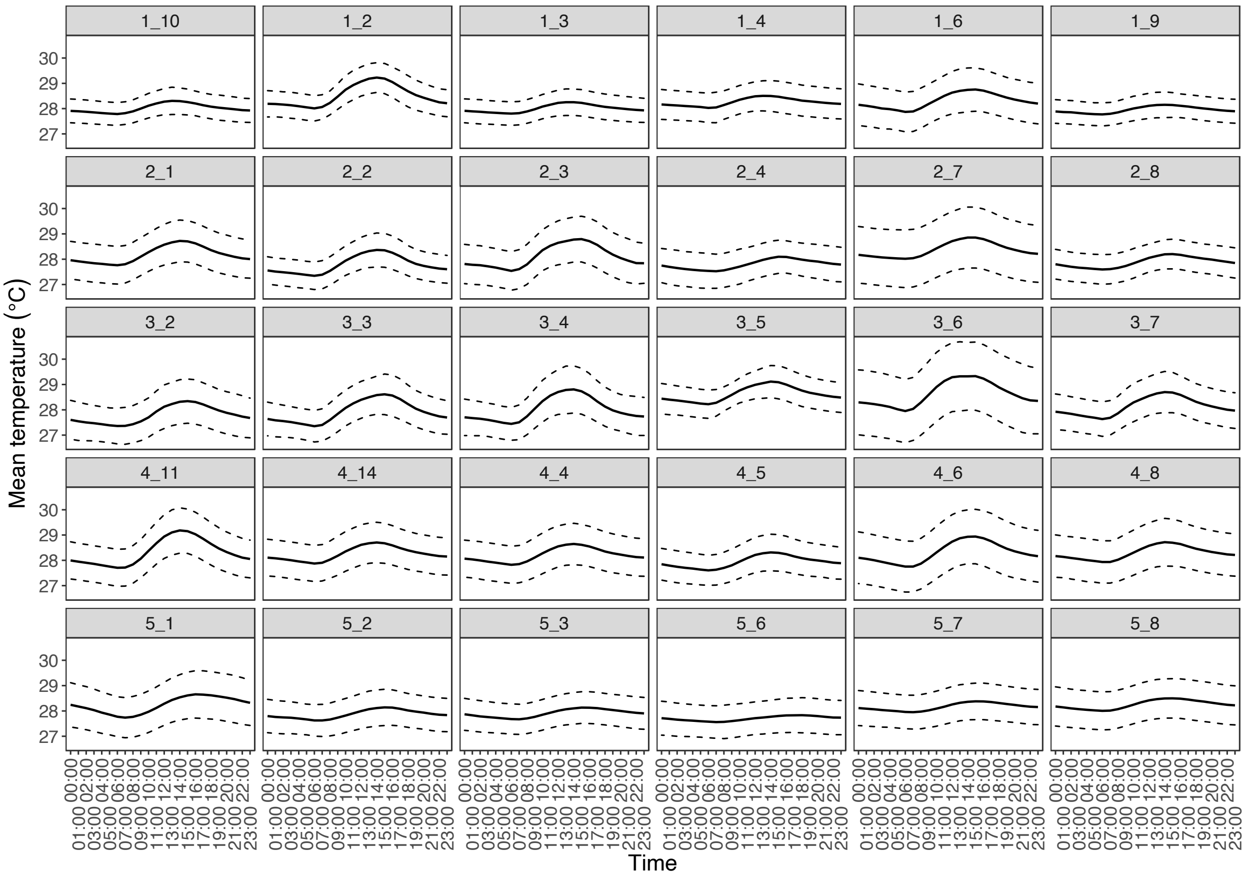


**Supplemental Figure 3 - Warm Season Temperature Variability**

Daily profiles of mean temperature (°C) per hour at each site, from 2017-2019 during the warm season (August 15th to October 15th). Dashed lines represent ± 1 standard deviation.

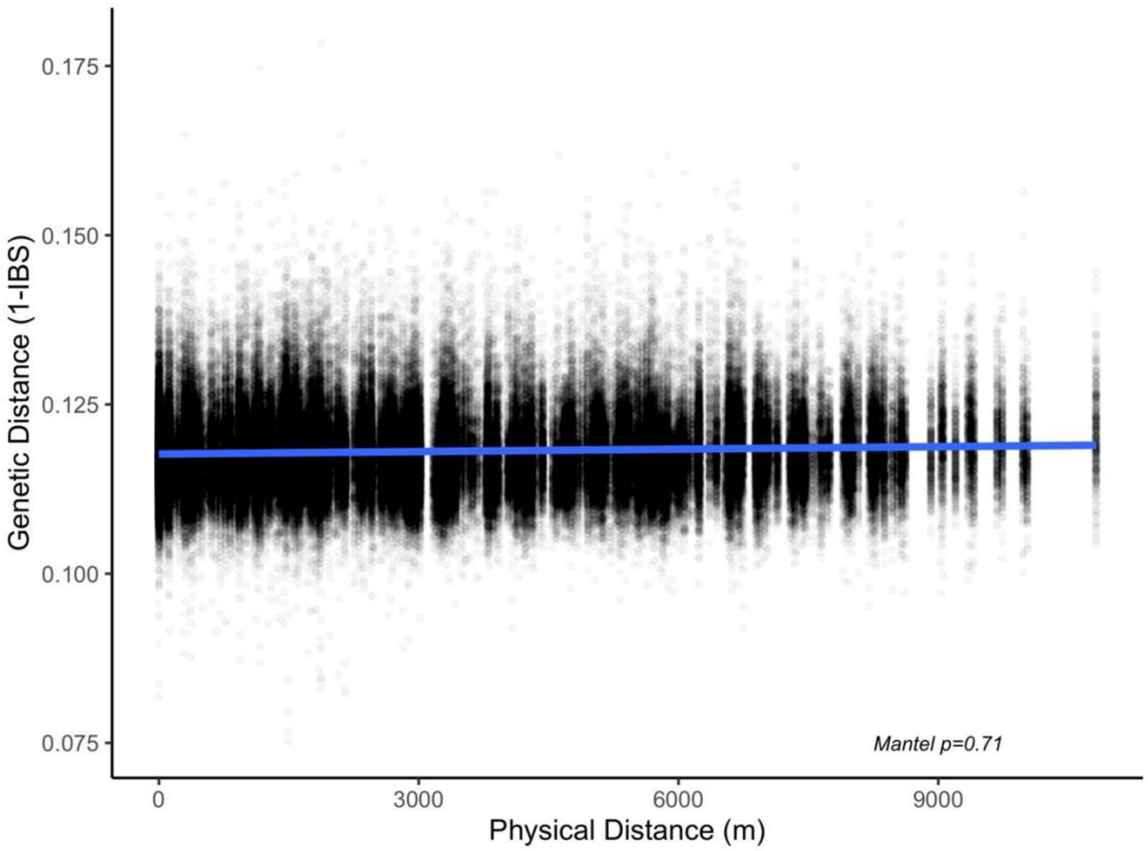


**Supplemental Figure 4 - Baywide Isolation by Distance**
Comparison of linear fit between genetic distance and physical distance. All samples at a site were considered to have the same coordinates and compared in pairwise fashion to all other samples.

**Supplemental Table 1 - Environmental Conditions for Each Site**
Summary of environmental conditions for each site. Summer calculations are data pooled from August 15 to October 15 for 2017-2019. Other studies column corresponds to closely overlapping site names from other studies.

**Supplemental Table 2 - Genetic Statistics for Each Site**
G:R is Genet-to-Ramet Ratio, equal to genotypes/samples where 1 equals no clonality. Mantel p is the (FDR corrected) significance of a mantel test comparing spatial and genetic distance within each site after randomly downsampling each genotype to a single sample in the case of clonality. Wilcox Distance is the (FDR corrected) p-value of a Wilcoxon test for spatial distance between samples of the same genotype and all corals at a site; NA values correspond to sites without any clones. Mean relatedness and mean relatedness without clones were calculated from genetic distance values derived from ANGSD.

**Supplemental Table 3 – dbRDA** PERMANOVA
Bray-Curtis dissimilarity summary of environmental drivers of genetic variance.
